# AncFlow: An Ancestral Sequence Reconstruction Approach for Determining Novel Protein Structural

**DOI:** 10.1101/2024.07.30.605920

**Authors:** Ryin Rouzbehani, Scott T. Kelley

## Abstract

The rapid growth of sequence data from high-throughput sequencing technologies has unveiled a vast number of previously unknown proteins, presenting a significant challenge in their functional characterization. Ancestral sequence reconstruction (ASR) has emerged as a powerful tool to elucidate the evolutionary history of protein families and identify sequence determinants of protein function. Here, we present AncFlow, an automated software pipeline that integrates phylogenetic analysis, subfamily identification, and ASR to generate ancestral protein sequences for structural prediction using state-of-the-art tools like AlphaFold. AncFlow streamlines the process of ASR by combining multiple sequence alignment, phylogenetic tree inference, subfamily identification, and ancestral sequence reconstruction from unaligned protein sequences. The reconstructed ancestral sequences are then subjected to structural prediction using AlphaFold, enabling the investigation of the structural basis of functional divergence within protein families. We validated AncFlow using two well-characterized protein families: acyltransferases and dehydrogenases. The pipeline successfully reconstructed ancestral sequences for multiple internal nodes of the phylogenetic trees, and their predicted structures were compared with those of extant proteins. By analyzing the structural similarities and differences between ancestral and extant proteins, we gained insights into the evolutionary mechanisms underpinning the functional diversification within these families. AncFlow demonstrates the potential of integrating ASR and structural prediction to unravel the structural basis of functional divergence in protein families. The insights gained from this approach can guide protein engineering efforts, facilitating the design of proteins with desired functions. As the amount of sequence data continues to grow, AncFlow provides a valuable tool for exploring the evolutionary landscape of proteins and accelerating the discovery of novel protein functions.

## INTRODUCTION

The rapid advancements in high-throughput sequencing technologies have revolutionized the field of genomics and proteomics, resulting in an exponential growth of sequence data. This wealth of sequence data has unveiled a plethora of previously unknown proteins, offering unprecedented insights into the molecular diversity of life [1]. However, characterizing unknown proteins remains a daunting challenge. Complete biochemical characterization of proteins is an expensive and time-consuming process, highlighting the need for efficient and innovative computational methods to model their structures and predict their functions [2].

Current approaches for predicting protein functions from sequence data encompass a range of computational methods. Sequence homology-based techniques, which involve comparing novel protein sequences to databases of functionally annotated proteins, have been widely employed [2]. Machine learning techniques, such as support vector machines and neural networks, have been applied to protein function prediction by leveraging datasets of proteins with known functions [3]. These methods can integrate various protein features, including sequence composition, physicochemical properties, and structural information, to enhance prediction accuracy. Nevertheless, the performance of these approaches is heavily dependent on the quality and comprehensiveness of the training data, which may be limited for novel protein families. Alternatively, another approach identifies the conserved functional domains found within protein sequences. Hidden Markov Models (HMMs) have proven particularly effective for identifying core functions of proteins [4]. By modeling the conserved sequence patterns and motifs associated with specific protein families and domains, HMMs can determine the potential functions for novel proteins based on their similarity to well-characterized protein families. However, the high functional diversity within protein families poses a significant challenge, as proteins with similar sequences may exhibit varying functions. For instance, bacterial porin proteins, which allow for the selective movement of hydrophilic solutes through the outer membrane of Gram-negative bacteria, can be classified into several subfamilies, such as ompC, ompF, phoE, nmpC, and ompN. Despite their sequence similarities, these subfamilies may have distinct functional properties and substrate specificities [5].

To address this issue, Ortiz-Velez et al. developed AutoPhy [6], an automated approach that combines phylogenetic analysis, dimensionality reduction, and clustering techniques to identify protein subfamilies with distinct functional roles. AutoPhy enables the detection of functionally diverse subfamilies within larger protein families, facilitating the discovery of novel protein functions. The vast array of proteins discovered through high-throughput sequencing has not only expanded our understanding of molecular diversity but has also opened new avenues for protein engineering. Protein engineering aims to design and create proteins with improved or distinct functions [7]. One promising approach in protein engineering is ancestral sequence reconstruction (ASR). ASR elucidates the sequences of ancient proteins based on the evolutionary relationships among modern-day proteins [8]. Reconstructing ancestral proteins can provide insights into the evolutionary history of protein families and helps identify sequence determinants of protein function. ASR has been successfully employed to engineer proteins with enhanced stability, catalytic activity, and altered substrate specificities [9]. For example, reconstructing ancestral versions of bacterial enzymes has led to the creation of highly stable and efficient biocatalysts that can withstand harsh industrial conditions [10]. Moreover, ASR has shed light on the evolutionary mechanisms underlying the functional diversification of protein families. By tracing the evolutionary trajectories of proteins, key mutations that have led to the emergence of new functions or the adaptation of proteins to different environmental conditions can be identified [11].

Here, we present AncFlow, a software pipeline designed to automate the various steps involved in producing ancestral sequences for use by structural prediction tools like AlphaFold [13]. Ancflow directly generates ASRs from protein families using unaligned protein sequences via the following steps: multiple sequence alignment, phylogenetic tree inference, subfamily identification, and ASR. The reconstructed ancestral sequences obtained from AncFlow were then further analyzed using AlphaFold to predict their three-dimensional structures. AncFlow was validated with the ASRs of multiple internal nodes from AutoPhy-resolved clades, and the respective structures of the internal nodes were then predicted with AlphaFold. The resultant structural predictions were then superimposed and aligned for comparative analysis, providing insights into the evolutionary mechanisms that have led to the diversification of the clade. Understanding these mechanisms offers valuable information about the design principles governing protein function, which can be adapted to engineer proteins for specific applications [16].

## METHODS

The workflow of the AncFlow pipeline outlined in Figure 1 begins with unaligned protein sequence data, and we tested the pipeline using data downloaded from the Pfam [4] and UniProt [19] databases. Multiple sequence alignments (MSAs) were performed on collections of sequences using MAFFT 7.49 [20], which leverages the Fast Fourier Transform (FFT) for rapid detection of homologous segments and utilizes a scoring system effective for sequences with large insertions/extensions and distantly related sequences. The MSA step is crucial for identifying conserved regions and exploring evolutionary relationships among sequences, which form the basis for subsequent phylogenetic and ancestral sequence analyses in AncFlow. The resulting alignments underwent an undetermined character check before proceeding to phylogenetic inference.

**Figure 1.**
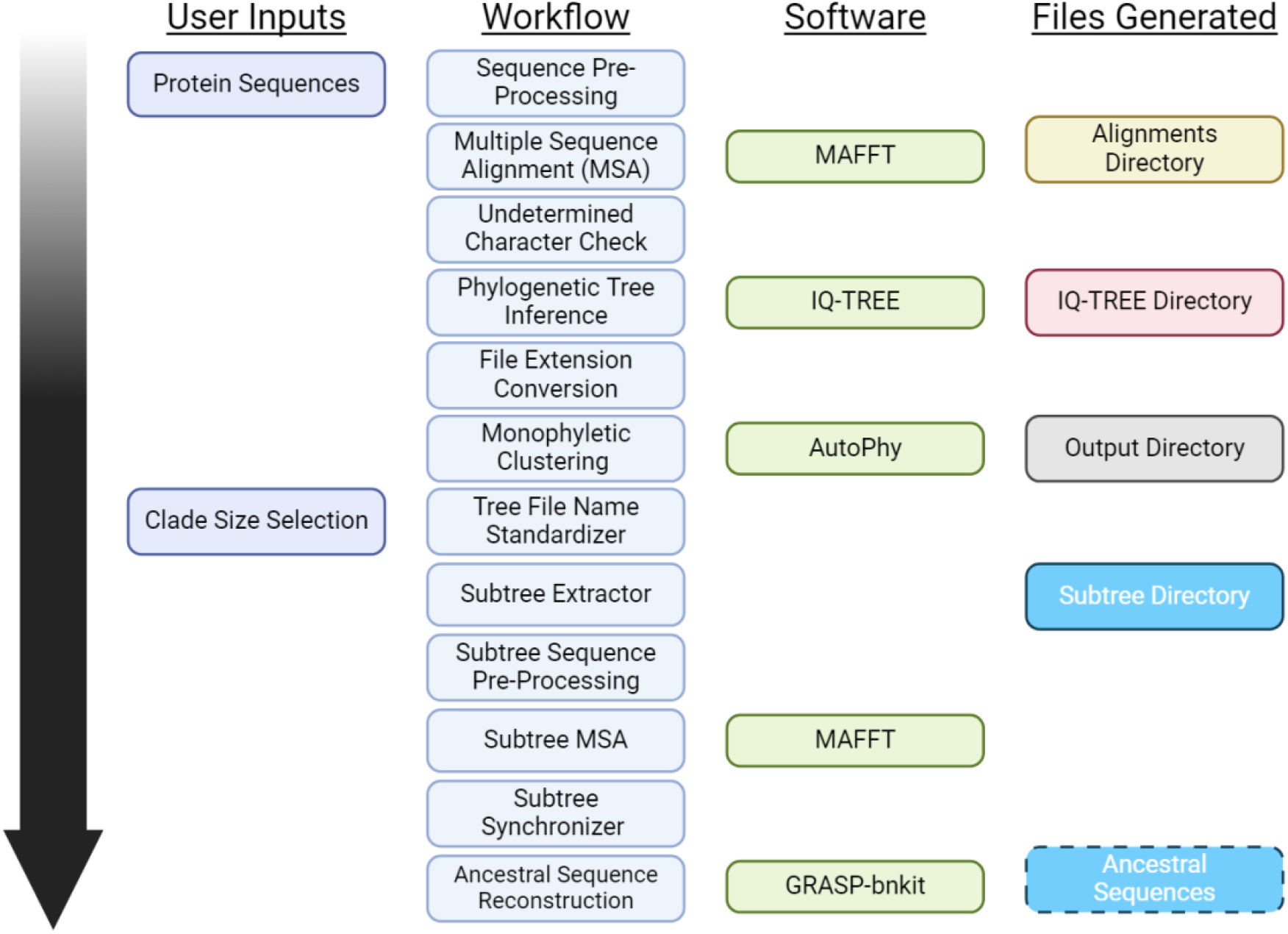
Workflow diagram of the AncFlow pipeline. User inputs include protein sequences and clade size selection. The workflow consists of sequence pre-processing, MSA, undetermined character check, phylogenetic tree inference, file extension conversion, monophyletic clustering, tree file name standardization, subtree extraction, subtree sequence pre-processing, subtree MSA using MAFFT, subtree synchronization, and ancestral sequence reconstruction employing GRASP-bnkit. The software used at each step is indicated, and the generated files, including alignment directories shown.

Phylogenetic inference of the MSAs was conducted using IQ-TREE 1.6.12 [21]. The inferred phylogenetic trees capture the evolutionary history of the protein families, providing a framework for identifying functionally distinct subfamilies and guiding the ancestral sequence reconstruction process in AncFlow. The resultant phylogenies were converted to the appropriate Newick file format and used as input for AutoPhy, resolving potential monophyletic subfamilies. Clades of interest were automatically extracted as independent phylogenetic subtrees based on user-defined clade size. These subtrees underwent sequence pre-processing, a final MSA, and synchronization before being subjected to ASR. Subsequently, ASR was performed on target nodes of subtrees using the Graphical Representation of Ancestral Sequence Predictions (GRASP) [12]. Finally, the predicted structures of the output ancestral sequences were achieved using AlphaFold and observed in ChimeraX [17].

### Sequence Datasets

To validate the AncFlow, two Swiss-Prot annotated protein family datasets were curated from the UniProt database. These families were chosen based on their extensive functional characterization and detailed annotations. The dataset comprises *acyltransferases* and *dehydrogenases*. To ensure compatibility with the AncFlow pipeline, the acquired protein sequences underwent a preprocessing step. The headers of the sequences were standardized to consistently indicate the origin of the sequences and truncated to include only the relevant information. This preprocessing aimed to improve consistency and compatibility across the datasets.

### Multiple Sequence Alignment

Following the preprocessing step, the standardized protein sequences were subjected to multiple sequence alignment using MAFFT. MAFFT is a widely used tool for generating high-quality alignments of multiple protein or nucleotide sequences. It employs various algorithms to optimize the alignment based on sequence similarity and evolutionary relationships. One of the algorithms used by MAFFT is FFT-NS-2, which is particularly effective for aligning numerous sequences. FFT-NS-2 utilizes the Fast Fourier Transform (FFT) to quickly identify homologous regions between sequences. It then performs a progressive alignment using a guide tree based on the identified homologies. This approach allows for efficient alignment of protein sequences while considering their evolutionary relationships. Another algorithm employed by MAFFT is L-INS-i, which is well-suited for aligning sequences with long insertions or extensions. L-INS-i iteratively refines the alignment by performing local pairwise alignments and incorporating consistency scores.

This iterative refinement process helps to improve the accuracy of the alignment, especially in regions with variable sequence lengths or low similarity. MAFFT also incorporates several protein-specific features to enhance the quality of the alignments. It utilizes amino acid substitution matrices, such as BLOSUM62, to assign appropriate scores for amino acid similarities and differences. Additionally, MAFFT considers the physicochemical properties of amino acids, such as hydrophobicity and size, to further refine the alignment. These protein-specific optimizations enable MAFFT to generate biologically meaningful alignments that accurately reflect the evolutionary relationships among the sequences. After obtaining the aligned sequences from MAFFT, AncFlow incorporates an additional processing step to handle ambiguous amino acid characters. In protein sequence alignments, the “X” character is commonly used to represent an unknown or ambiguous amino acid.

However, downstream tools do not recognize and handle the “X” character appropriately. To address this issue, a sequence processing function was implemented to replace all occurrences of “X” with a gap character “-”. This replacement step ensures that the aligned sequences are properly formatted and can be seamlessly used in subsequent analyses.

### Phylogenetic Analysis

Phylogenetic inference of the MSAs was achieved with IQ-TREE, a maximum likelihood-based tool, using an LG+G+F substitution model. The LG model, developed by Le and Gascuel [22], is an amino acid substitution matrix specifically designed for protein sequence analysis. It captures the evolutionary patterns and substitution rates observed in a wide range of protein families. The +G parameter accounts for rate heterogeneity across sites, allowing for varying evolutionary rates among different positions in the sequence alignment. The +F parameter incorporates empirical amino acid frequencies, ensuring that the model reflects the observed amino acid composition of the sequences. To assess the statistical support for the phylogenetic relationships, 1000 bootstrap replicates were performed. IQ-TREE constructs a phylogenetic tree for each bootstrap replicate and calculates the proportion of replicates supporting each branch of the consensus tree. IQ-TREE employs a combination of nearest neighbor interchange (NNI) and subtree pruning and regrafting (SPR) algorithms to search for the optimal tree topology. These algorithms explore the space of possible tree topologies by making local rearrangements and evaluating the likelihood of each configuration. The software automatically parallelizes the tree inference process, optimizing computational resources based on the available hardware. The inferred phylogenetic trees, along with statistical information and branch support values, are saved in the “iqtree” directory.

### Clustering and Protein Subfamily Identification

The resulting phylogenetic trees output by IQ-TREE underwent clustering and subfamily identification by AutoPhy. AutoPhy leverages the evolutionary information encoded in the tree topology and branch lengths to delineate functionally distinct subgroups. AutoPhy beings by extracting the pairwise evolutionary distances using the DendroPy library [23], and then creating a pairwise distance matrix (PDM) which serves as the input for subsequent dimensionality reduction. To mitigate noise while preserving the intrinsic data structure, AutoPhy employs Uniform Manifold Approximation and Projection (UMAP) [24], a non-linear technique that maps the high-dimensional PDM to a lower-dimensional space. The reduced matrix is then partitioned into putative subfamilies using a Gaussian mixture model optimized via the Expectation-Maximization algorithm (GMMEM). The Bayesian Information Criterion (BIC) is employed to objectively select the model with the most appropriate number of clusters. To ensure that the resulting clusters correspond to monophyletic groups in the original tree, AutoPhy recursively refines non-monophyletic clusters using the “monophyletic” function in Biopython [25] until all subfamilies represent coherent evolutionary units. The identification of monophyletic subfamilies is crucial for the AncFlow pipeline, as it ensures that the subsequent ancestral sequence reconstruction and functional analyses are performed on evolutionarily meaningful subgroups. As a final step, AutoPhy derives a log ratio score for each subfamily by contrasting its internal divergence (measured as the mean pairwise distance) against the length of its subtending branch, providing insights into the relative evolutionary rates across subfamilies and potentially highlighting clades subjected to distinctive selective pressures. Upon successful completion, the “output” directory is populated with the clustered phylogenetic tree, the UMAP projection visualizing the reduced-dimensional representation of the pairwise distance matrix, and the Gaussian mixture model plot depicting the fitted components used for clustering.

### Subtree Extraction

Following the identification of monophyletic subfamilies by AutoPhy, a Python script extracts subtrees corresponding to each subfamily from the clustered phylogenetic tree. The subtree extraction step focuses the AncFlow analysis on specific subgroups of interest, reducing computational complexity and allowing for more accurate ancestral sequence reconstruction within the targeted subfamilies. The script prompts the user to specify a minimum clade size threshold, ensuring that only subtrees with a sufficient number of sequences are considered for further analysis. During the parsing process, the script identifies the taxon labels associated with each sequence in the subtree, which typically follow a standardized format such as “sp_accession|clade_number:branch_length”. The script extracts the accession numbers and clade numbers from these labels, allowing for the retrieval of corresponding protein sequences from the NCBI GenBank database using the Biopython library. The retrieved sequences and extracted subtrees are then saved in FASTA and Newick formats, respectively, in a designated output directory with file names reflecting the corresponding clade number. After the subtree extraction process, another Python script standardizes the sequence headers in the FASTA files to the UniProt format, which follows the convention “>tr|accession”. The script recursively searches the designated directory for FASTA files, processes the sequences, and modifies the headers using a dedicated function that extracts the accession number from the original FASTA header and constructs the UniProt-compliant header. The modified sequences are initially written to temporary files and then replace the original FASTA files upon successful completion of the header conversion process. The AncFlow pipeline then performs another MSA on the extracted subtree sequences using MAFFT. This additional MSA step refines the alignment within each subfamily, enabling more accurate ancestral sequence reconstruction by focusing on the evolutionary relationships and sequence diversity specific to each subgroup. To ensure consistency and enable accurate ancestral sequence reconstruction, a Python script synchronizes the sample names between the aligned FASTA files and their corresponding Newick tree files. The script extracts the sample names from the Newick tree file, and constructs a dictionary mapping the accession numbers to the extracted sample names. The script then processes the corresponding FASTA file, replacing the original sample names in the header lines with the sample names obtained from the Newick tree file, using the accession numbers as the matching key. The resulting subtrees and their corresponding sequence files, with standardized headers and synchronized sample names, are advanced to ASR.

### Ancestral Sequence Reconstruction

Following the synchronization of subtree sequences and their corresponding phylogenetic trees, ASR was performed using the GRASP tool. GRASP employs a maximum likelihood approach to approximate ancestral sequences, leveraging the evolutionary information encoded in the phylogenetic trees and the sequence diversity present in the aligned protein sequences. GRASP utilizes partial order graphs (POGs) to represent sequence content, including insertions and deletions, enabling the handling and visualization of indel events across ancestral sequences. POGs offer a flexible and efficient way to capture the complex evolutionary relationships among sequences, allowing for the representation of multiple possible ancestral states and the handling of alignment ambiguities [12]. By considering the evolutionary relationships and the observed sequence variations, GRASP reconstructs the most probable ancestral sequences at the target nodes of the subtrees. The reconstruction process in GRASP involves three steps. First, the history of indel events is inferred, determining the presence or absence of specific indels at each ancestral node. Second, GRASP assigns the most likely amino acid characters at each ancestral position based on indel history and the observed sequence variations. This is achieved through a joint reconstruction approach that considers the entire evolutionary context. Third, GRASP constructs ancestor POGs that represent the approximated sequence content, capturing the ancestral states and the potential ambiguities arising from multiple equally probable scenarios [12]. Finally, the completed ancestral sequences of all subtree internal nodes are resolved and prepared for structural prediction.

### Protein Structure Prediction

The reconstructed ASR were subjected to protein structure prediction using AlphaFold 2.3.2, a modern deep learning-based method that predicts protein structures from primary sequences. AlphaFold leverages evolutionary information derived from multiple sequence alignments, physical principles such as energy minimization, and advanced machine learning techniques, including attention-based neural networks, to generate high-quality 3D models of protein structures. The localcolabfold 1.5.5 [26] implementation of AlphaFold was employed to predict the structures of the ancestral sequences. Localcolabfold uses the AlphaFold model for protein structure prediction without the need for extensive cloud based computational resources. The ancestral sequences were provided as input to localcolabfold, which generated predicted 3D structures in PDB format, along with per-residue confidence scores (pLDDT) visualized in ChimeraX. The predicted structures of the ancestral proteins and their respective extant descendants were superimposed and analyzed using MatchMaker [18] to elucidate key conserved and divergent structural motifs responsible for functional variance.

## RESULTS AND DISCUSSION

AncFlow was tested on a 2019 MacBook Pro with 32GB of 2667 MHz DDR4 RAM and 2.6 GHz 6-Core Intel Core i7. AncFlow successfully analyzed 173 sequences from the acyltransferase protein family in approximately 2 hours and 35 minutes and 213 sequences from the dehydrogenase protein family in approximately 2 hours and 18 minutes. The multiple sequence alignment (MSA) step took approximately 1 minute and 32 seconds for the acyltransferase dataset and 35 seconds for the dehydrogenase dataset. Phylogenetic tree inference was the most time-consuming step, requiring 2 hours and 28 minutes for acyltransferases and 2 hours and 18 minutes for dehydrogenases (Figure 2). Autophy identified 51 and 62 putative clusters in the acyltransferase and dehydrogenase phylogenetic trees, respectively. The subtree extraction step processed 31 clades for acyltransferases and 38 clades for dehydrogenases, with a minimum clade size of 3 sequences. ASR successfully reconstructed ancestral sequences for all extracted subtrees in both protein families. Nodes belonging to subtrees exhibiting high bootstrap values and potential within-group differentiation were isolated for structural prediction.

**Figure 2.**
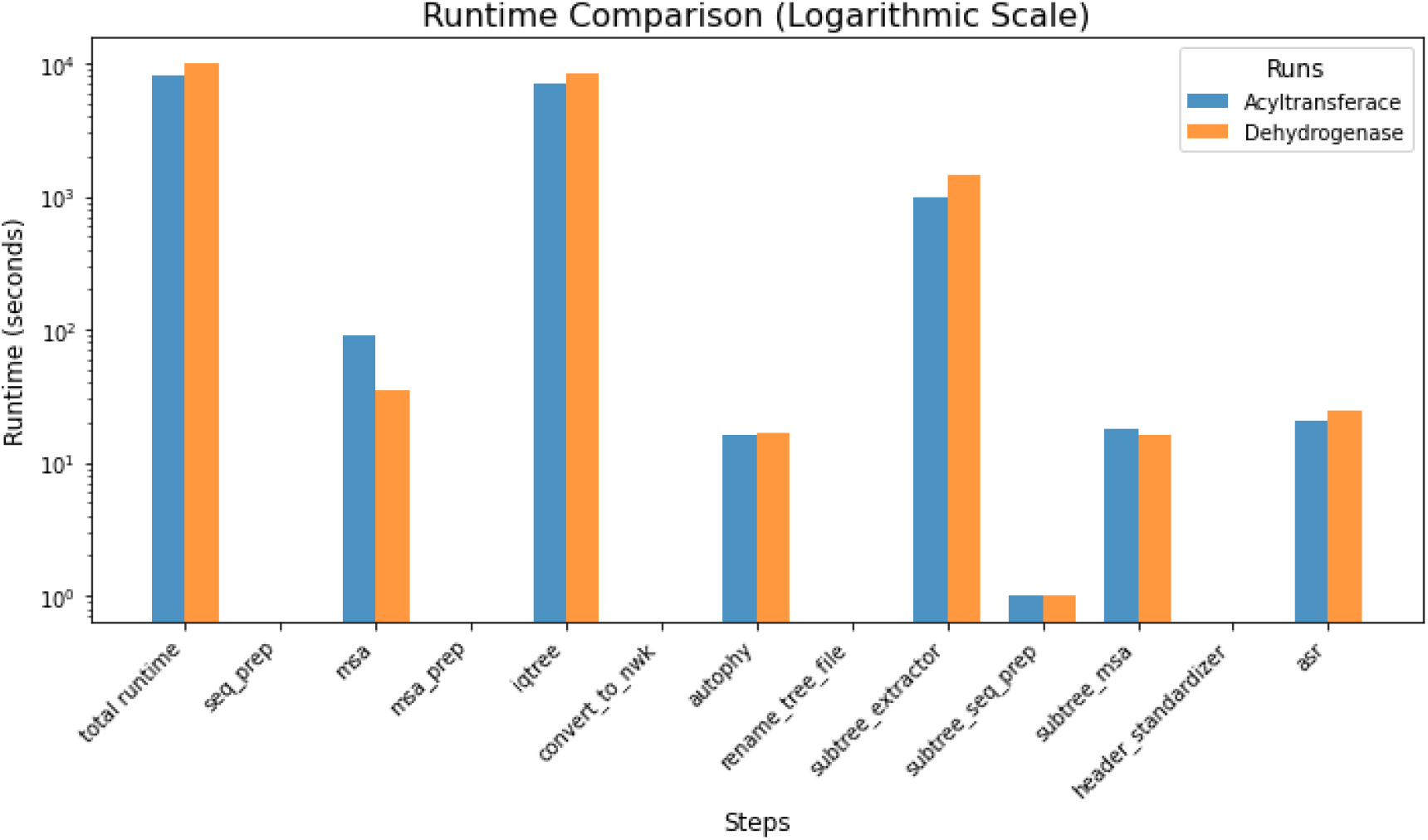
Runtime breakdown of the AncFlow pipeline steps for the acyltransferase and dehydrogenase protein families, shown on a logarithmic scale. Phylogenetic tree inference (IQ-TREE) dominates the total runtime, taking hours to complete. In contrast, the other steps are significantly faster, requiring minutes or less.

To optimize performance and resource utilization, AncFlow was executed with 2 cores, to prevent resource contention and performance degradation caused by over-parallelization. This allowed the pipeline to efficiently utilize computational resources and minimize the overall runtime. IQ-TREE, used for phylogenetic tree inference, also employed multi-threading to enhance performance. The optimal number of threads was determined to be 9 for the acyltransferase dataset and 6 for the dehydrogenase dataset. In terms of memory usage, IQ-TREE required approximately 322 MB of RAM for the acyltransferase family and 314 MB of RAM for the dehydrogenase family. It’s important to note that memory usage and CPU utilization may vary depending on the specific hardware configuration and the size and complexity of the input datasets.

### Acyltransferases Protein Family

Acyltransferases are a diverse family of enzymes that catalyze the transfer of an acyl group from a donor molecule to an acceptor substrate, playing crucial roles in various biological processes such as lipid biosynthesis, cell signaling, and metabolism. In this study, the main catalytic activity of each extant sequence was assigned as a functional marker to investigate the evolutionary history and potential functional divergence within a subtree of the acyltransferase protein family (Figure 3). By comparing the catalytic activities and predicted protein structures of the extant sequences and their ancestral counterparts, insights were gained into the structural basis of functional evolution in this diverse family of enzymes.

**Figure 3.**
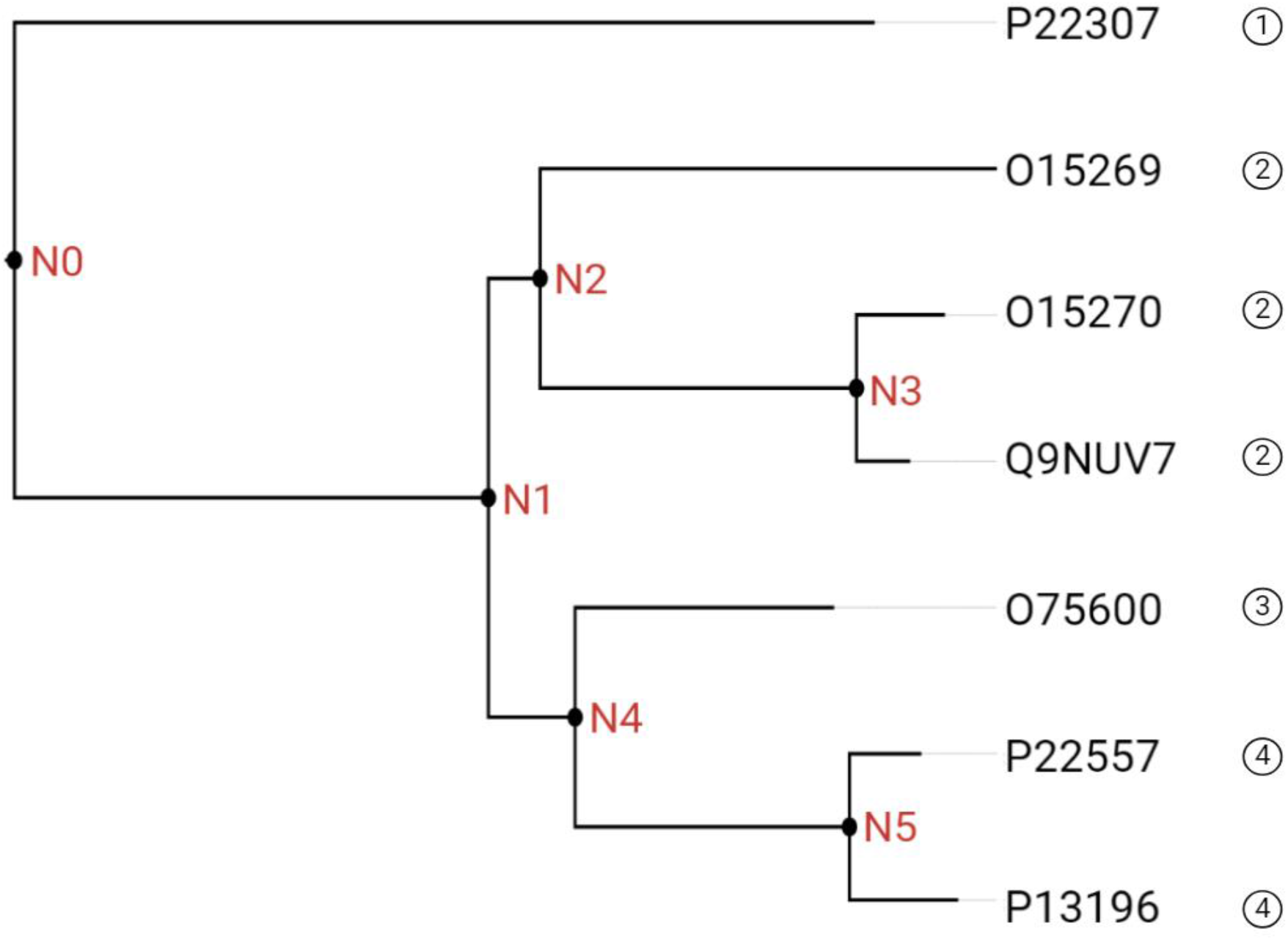
Acyltransferace protein family subtree with ancestral nodes labeled N0 to N5. The tree branches from the root node N0, splits into two main clades at N1, and further diversifies at nodes N2, N3, N4, and N5. The terminal nodes represent extant sequences, labeled with their respective accession numbers: P22307, O15269, O15270, Q9NUV7, O75600, P22557, and P13196. The circled numbers at the tips of the tree indicate enzymatic reactions shown in Table 1.

**Table 1.** Primary known or predicted catalytic activities of the extant sequences in the selected acyltransferase subtree.

| #* | Primary Catalytic Activity |
| --- | --- |
| 1 | Choloyl-CoA + Propanoyl-CoA → 3α,7α,12α-trihydroxy-24-oxo-5β-cholestan-26-oyl-CoA + CoA |
| 2 | H<sup>+</sup> + Hexadecanoyl-CoA + L-serine → 3-oxosphinganine + CO<sub>2</sub> + CoA |
| 3 | Acetyl-CoA + Glycine → (2S)-2-amino-3-oxobutanoate + CoA |
| 4 | Glycine + H<sup>+</sup> + Succinyl-CoA → 5-aminolevulinate + CO<sub>2</sub> + CoA |

\* Refers to number at tips of phylogenetic tree in Figure 3.

The extant sequences in the selected subtree exhibited distinct catalytic activities, reflecting the functional diversity within the acyltransferase family. P13196 and P22557, both annotated as 5-aminolevulinate synthases, catalyze the pyridoxal 5’-phosphate (PLP)-dependent condensation of succinyl-CoA and glycine to form aminolevulinic acid (ALA), with CoA and CO2 as by-products. O75600, a 2-amino-3-ketobutyrate coenzyme A ligase, catalyzes the cleavage of 2-amino-3-oxobutanoate to glycine and acetyl-CoA. O15270, Q9NUV7, and O15269, all identified as serine palmitoyltransferases, catalyze the initial and rate-limiting step in sphingolipid biosynthesis by condensing L-serine and activated acyl-CoA to form long-chain bases. Lastly, P22307, a sterol carrier protein, plays a crucial role in the peroxisomal oxidation of branched-chain fatty acids and catalyzes the last step of the peroxisomal beta-oxidation of branched chain fatty acids and the side chain of bile acid intermediates.

To investigate the structural basis of these functional differences, the protein structures of the extant sequences were predicted using AlphaFold (Figure 4). However, structural variations were evident between the extant sequences, particularly in the arrangement and orientation of the secondary structure elements. These structural differences likely reflect the functional diversity observed in the catalytic activities of the extant sequences. The ancestral sequences at the internal nodes of the subtree (N0 to N5) were reconstructed and their protein structures were predicted using AlphaFold (Figure 5). The predicted ancestral structures exhibited notable variations, particularly in the arrangement and orientation of the secondary structure elements, suggesting potential functional divergence throughout the evolutionary history of the family.

**Figure 4.**
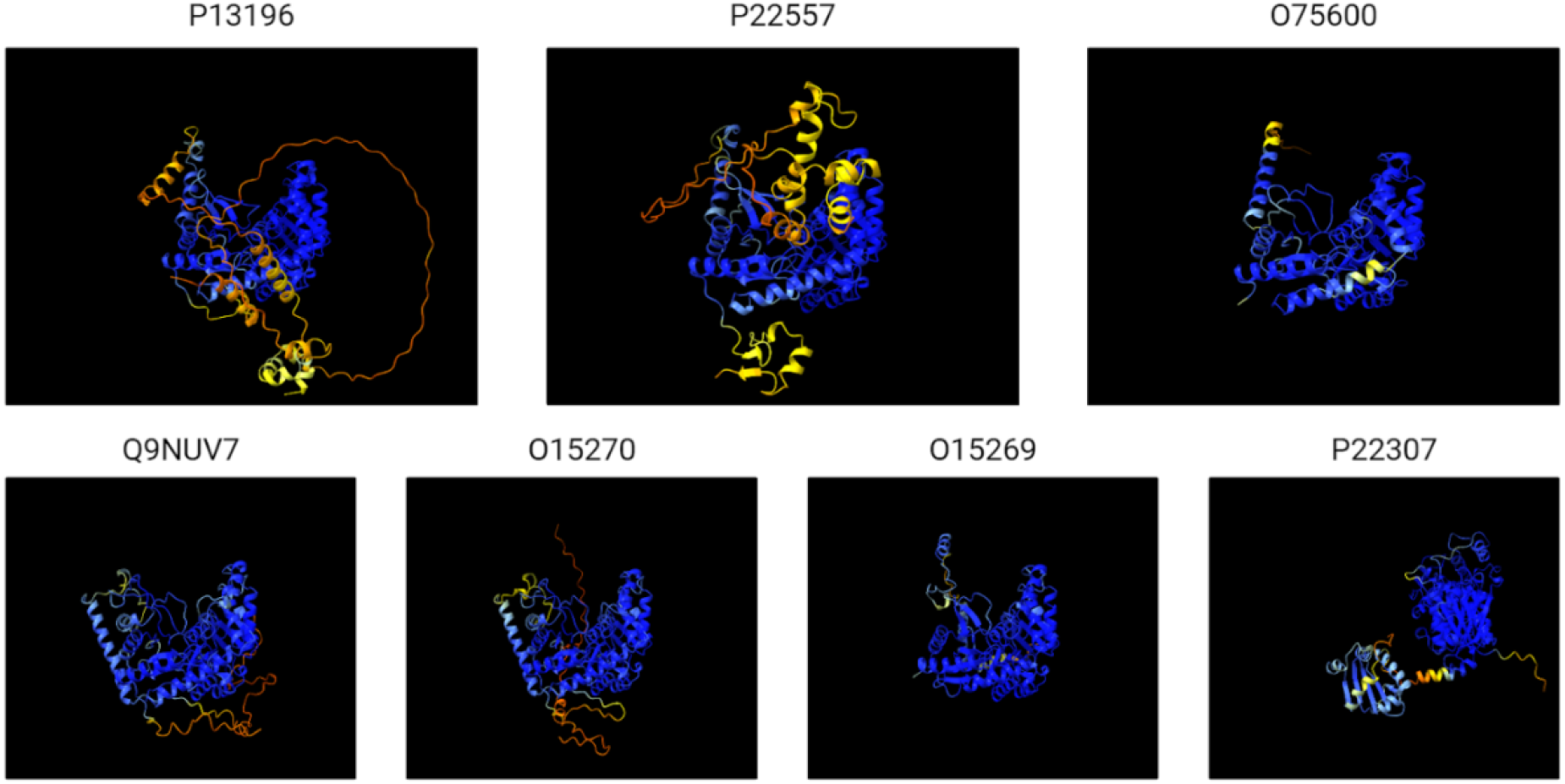
Predicted protein structures for extant sequences P13196, P22557, O75600, Q9NUV7, O15270, O15269, and P22307, generated using AlphaFold. Structural variations between the extant sequences are evident, particularly in the arrangement and orientation of the secondary structure elements.

**Figure 5.**
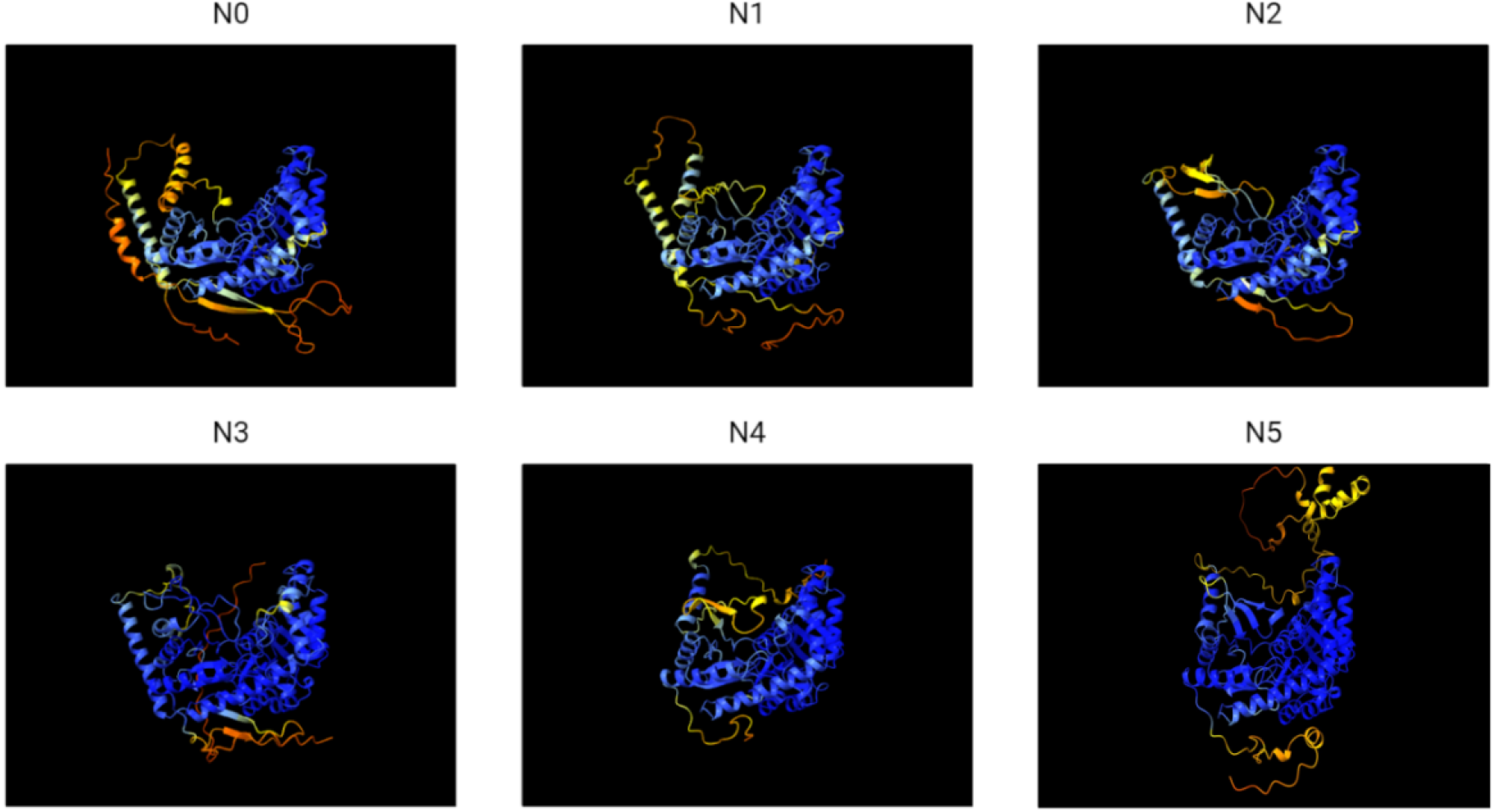
Predicted protein structures for ancestral nodes N0 to N5, generated using AlphaFold. Structural variations between the ancestral nodes are evident, particularly in the arrangement and orientation of the secondary structure elements, suggesting potential functional divergence.

By comparing the predicted structures of the ancestral nodes with those of the extant sequences, structural similarities and differences were observed that could hint at the functional evolution of the catalytic activities. For example, the predicted structures of N5 and its descendant extant sequences, P13196 and P22557, exhibited similar structural features, consistent with their shared catalytic activity as 5-aminolevulinate synthases.

Interestingly, the predicted structure of N4 also displayed great similarity to its descendant extant sequences, O75600, P22557, and P13196, which have distinct main catalytic activities. These structural differences suggest potential functional divergence at this evolutionary juncture [27]. Similarly, the predicted structures of N3 and its descendant extant sequences, O15270 and Q9NUV7 showed structural similarities, consistent with their shared function as serine palmitoyltransferases.

### Dehydrogenase Protein Family

The dehydrogenase protein family was investigated using the main catalytic activity of each extant sequence as a functional marker to explore the evolutionary history and potential functional divergence within the family (Figure 6). By comparing the catalytic activities and predicted protein structures of the extant sequences and their ancestral counterparts, insights were gained into the structural basis of functional evolution in this diverse family of enzymes.

**Figure 6.**
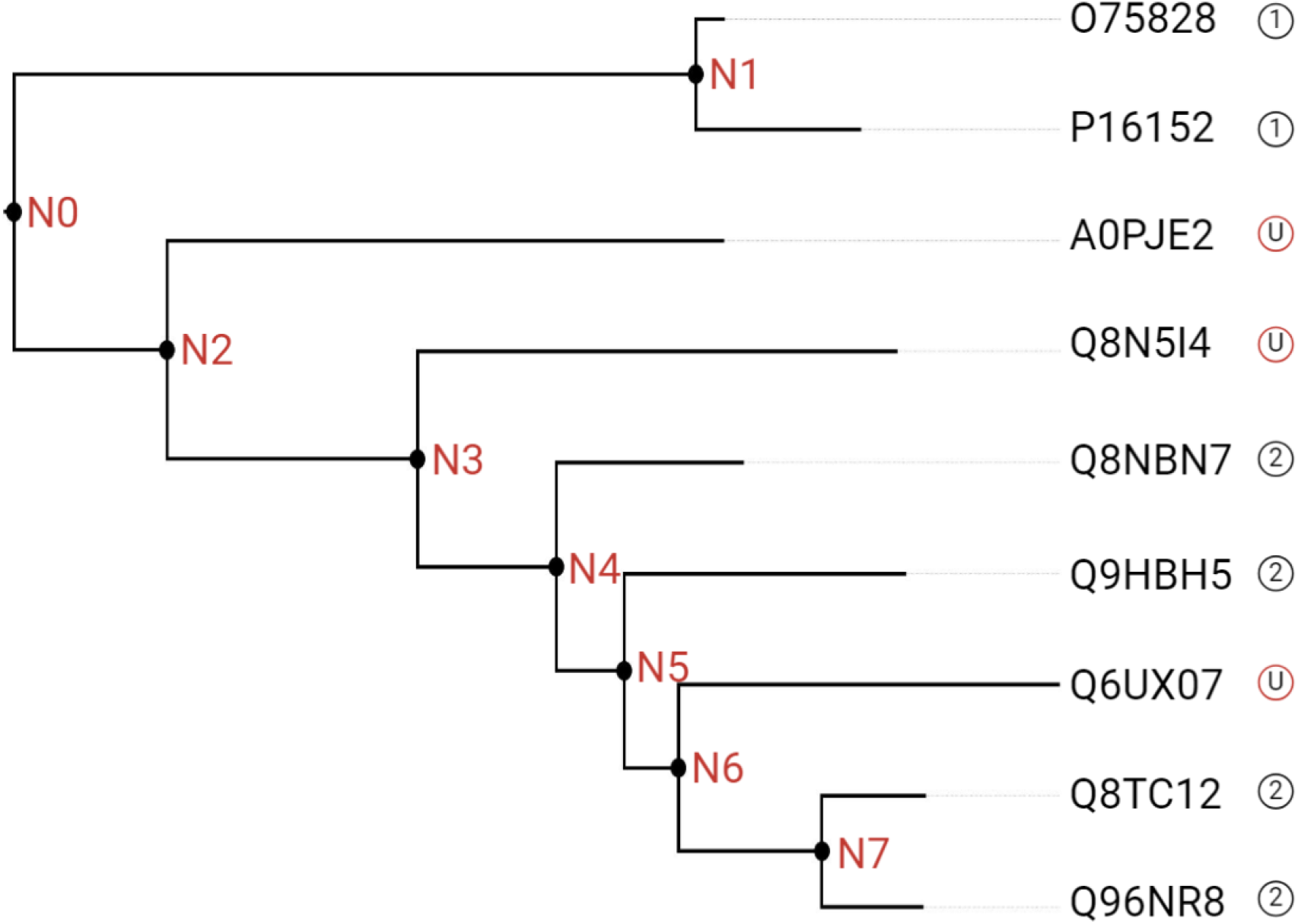
Dehydrogenase protein family subtree, with ancestral nodes labeled N0 to N7. The tree branches from the root node N0, splits into two main clades at N2, and further diversifies at subsequent nodes. The terminal nodes represent extant sequences, labeled with their respective UniProt accession numbers: O75828, P16152, A0PJE2, O8N5J4, Q8NBN7, Q9HBH5, Q6UX07, Q8TC12, and Q96NR8. The circled numbers at the tips of the tree indicate enzymatic reactions shown in Table 2. “U” denotes unknown catalytic activity.

**Table 2.** Primary known or predicted catalytic activities of the extant sequences in the selected dehydrogenase subtree.

| #* | Primary Catalytic Activity |
| --- | --- |
| 1 | A secondary alcohol + NADP<sup>+</sup> = A Ketone + H<sup>+</sup> + NADPH |
| 2 | All-trans-retinol + NADP<sup>+</sup> = All-trans-retinal + H<sup>+</sup> + NADPH |

\* Refers to number at tips of phylogenetic tree in Figure 6.

The extant sequences in the selected subtree exhibited distinct catalytic activities. O75828 and P16152, annotated as carbonyl reductases, catalyze the NADPH-dependent reduction of carbonyl compounds, including quinones, prostaglandins, and various xenobiotics. A0PJE2 and Q8N5I4 are putative oxidoreductases with unknown catalytic activities. Q8NBN7, Q9HBH5, Q8TC12, and Q96NR8 are retinol dehydrogenases that oxidize retinol to retinal using NADP+ as a cofactor, with varying substrate preferences for different retinol isomers. Q6UX07 is another putative oxidoreductase with unknown catalytic activity.

To investigate the structural basis of these functional differences, the protein structures of the extant sequences were predicted using AlphaFold (Figure 7). The predicted structures exhibited a Rossmann-like fold, characteristic of NAD(P)-binding proteins, with alternating alpha-helices and beta-strands forming a central beta-sheet. However, structural variations were evident between the extant sequences, particularly in the arrangement and orientation of the secondary structure elements, likely reflecting the functional diversity observed in their catalytic activities.

**Figure 7.**
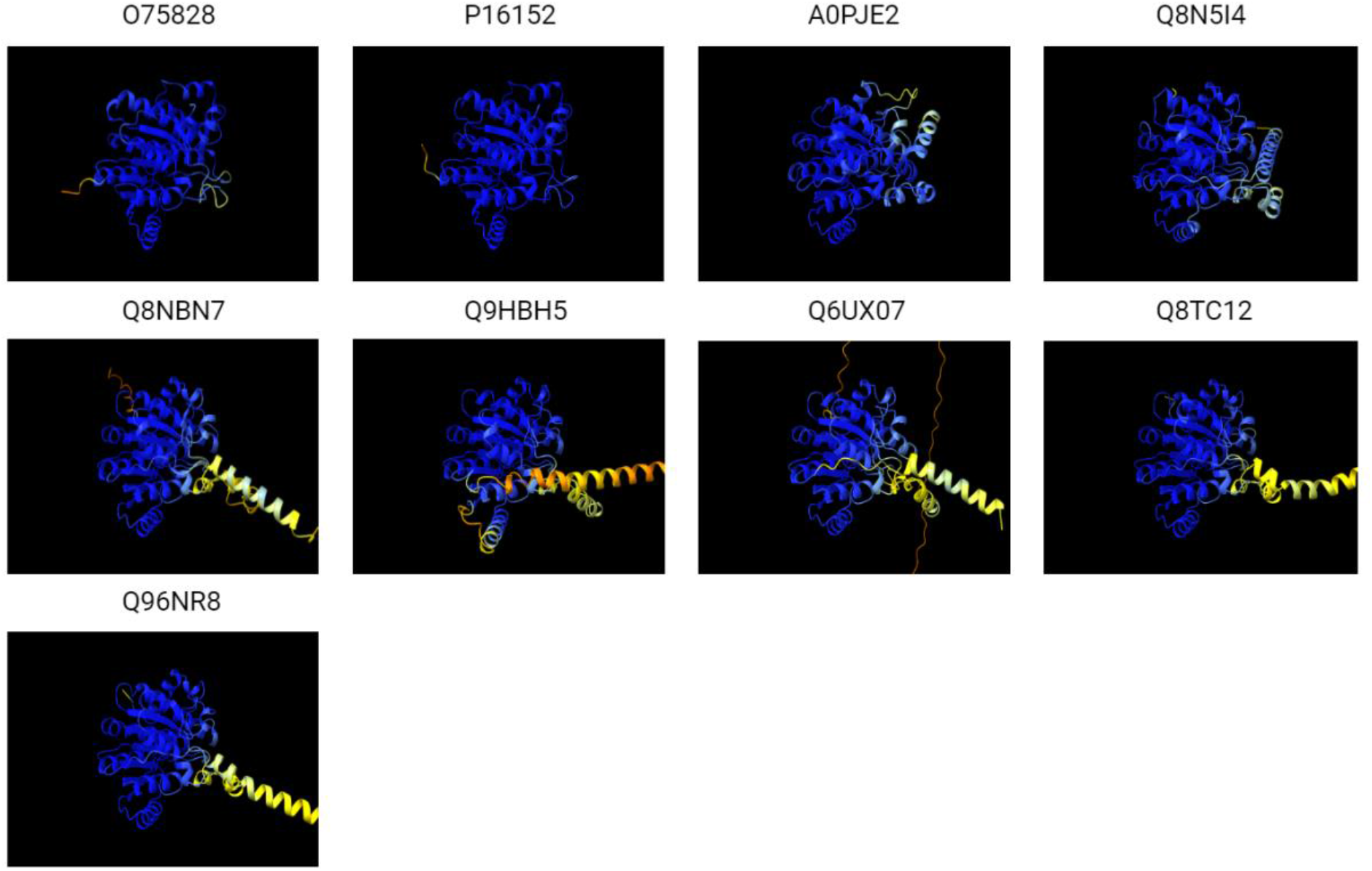
Predicted protein structures for extant sequences P13196, P22557, O75600, Q9NUV7, O15270, O15269, and P22307, generated using AlphaFold.

To gain insights into the evolutionary history of the dehydrogenase family, the ancestral sequences at the internal nodes of the subtree (N0 to N7) were reconstructed, and their protein structures were predicted using AlphaFold (Figure 8). The predicted ancestral structures also exhibited notable variations, particularly in the arrangement and orientation of the secondary structure elements, suggesting potential functional divergence throughout the evolutionary history of the family. By comparing the predicted structures of the ancestral nodes with those of the extant sequences, structural similarities and differences were observed that could hint at the functional evolution of the catalytic activities [28]. For example, the predicted structures of N7 and its descendant extant sequences, Q8TC12 and Q96NR8, exhibited similar structural features, consistent with their shared function as retinol dehydrogenases with a preference for NADP+ and activity towards various retinol isomers. This structural similarity suggests that the ancestral protein at N7 likely possessed a similar retinol dehydrogenase activity. In contrast, the predicted structure of N6 displayed notable differences compared to its descendant extant sequence, Q6UX07, which has an unknown catalytic activity. These structural differences suggest potential functional divergence at this evolutionary juncture, with the ancestral protein at N6 possibly having a different catalytic activity than its descendant. The predicted structures of N1 and its descendant extant sequences, O75828 and P16152, showed structural similarities, consistent with their shared function as carbonyl reductases. This structural similarity suggests that the ancestral protein at N1 likely possessed a similar carbonyl reductase activity, catalyzing the NADPH-dependent reduction of carbonyl compounds. However, the predicted structures of N2 and N3 exhibited differences compared to their descendant extant sequences, A0PJE2, Q8N5I4, Q8NBN7, and Q9HBH5, which have unknown or diverse catalytic activities. These structural variations indicate potential functional divergence at these evolutionary points, with the ancestral proteins at N2 and N3 possibly having different catalytic activities than their descendants.

**Figure 8.**
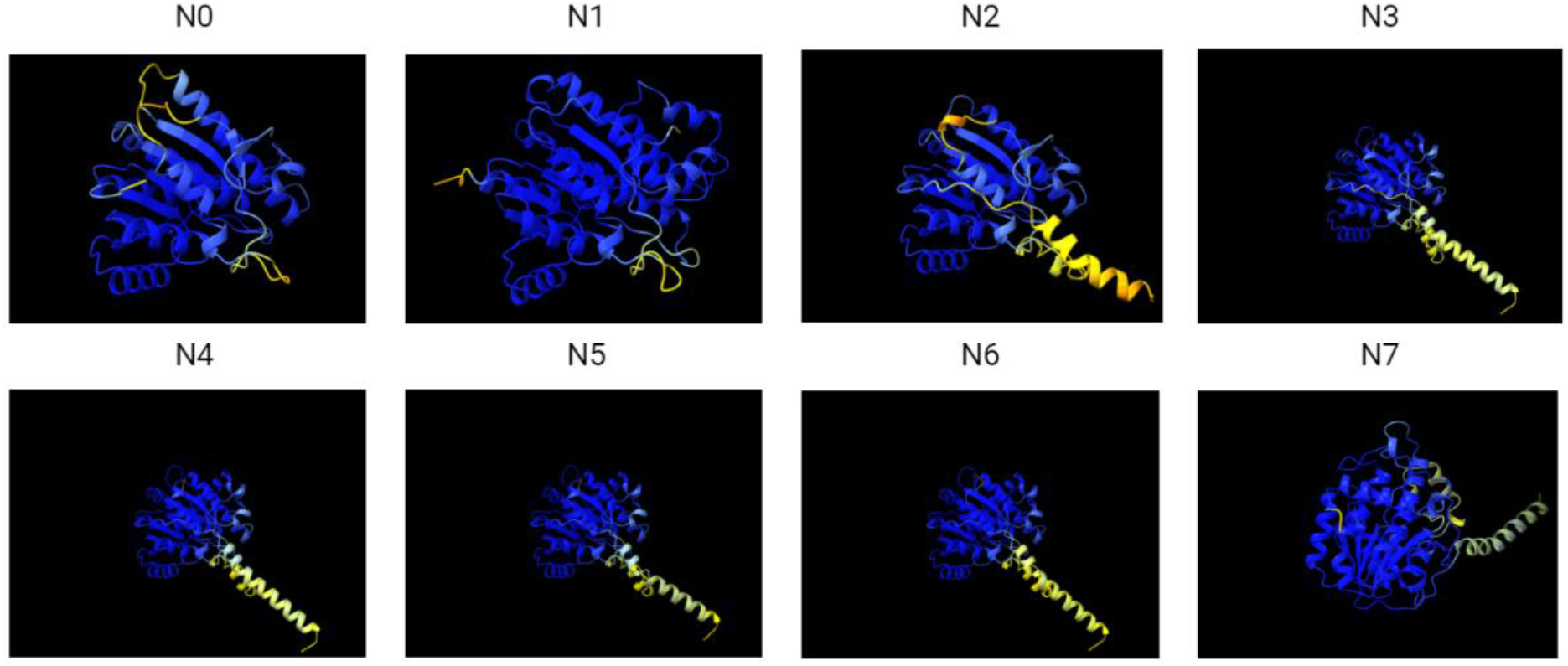
Predicted protein structures for ancestral nodes N0 to N5, generated using AlphaFold.

## CONCLUSION

By integrating phylogenetic analysis, subfamily identification, and ancestral sequence reconstruction, AncFlow provides the foundation needed for further exploration of structural functional divergence within protein families by structural prediction tools. The insights gained from AncFlow can guide further experimental studies and contribute to our understanding of the evolutionary mechanisms shaping the functional diversity of proteins [29]. The reconstructed ancestral sequences and their predicted structures can serve as valuable starting points for protein engineering efforts, potentially leading to the development of novel enzymes with desired properties for biotechnological applications.

## Data and Code Availability

All the code, including test files and installation instructions, can be found at https://github.com/rrouz/AncFlow.

## Acknowledgements

We thank J. Love and J. Sukumaran for helpful comments on the project and manuscript. We also think members of the Kelley Lab for testing out AncFlow.

## REFERENCES

[1] E. Levy Karin, R. Hershberg, and M. Barak, “The dark matter of the human proteome: Insights into the prevalence, characteristics, and functional implications of proteins of unknown function,” Proteomics, vol. 21, no. 3–4, p. 2000238, 2021.

[2] A. Dalkiran et al., “ECPred: a tool for the prediction of the enzymatic functions of protein sequences based on the EC nomenclature,” BMC Bioinformatics, vol. 19, no. 1, pp. 1–13, 2018.

[3] R. Fa, D. Cozzetto, C. Wan, and D. T. Jones, “Predicting human protein function with multi-task deep neural networks,” PLoS One, vol. 13, no. 6, p. e0198216, 2018.

[4] S. El-Gebali et al., “The Pfam protein families database in 2019,” Nucleic Acids Res., vol. 47, no. D1, pp. D427–D432, 2019.

[5] T. X. Nguyen, E. R. Alegre, and S. T. Kelley, “Phylogenetic Analysis of General Bacterial Porins: A Phylogenomic Case Study,” J. Mol. Microbiol. Biotechnol., vol. 11, no. 6, pp. 291–301, Nov. 2006.

[6] A. N. Ortiz-Velez, J. Sukumaran, R. Rouzbehani, and S. T. Kelley, “AutoPhy: Automated phylogenetic identification of novel protein subfamilies,” PLoS One, vol. 19, no. 1, p. e0291801, 2024.

[7] R. Borne, A. D. Levi, and A. R. Fersht, “Protein engineering: from modelling and design to biotechnological applications,” Curr. Opin. Biotechnol., vol. 76, p. 102723, 2022.

[8] Y. Gumulya et al., “Engineering highly functional thermostable proteins using ancestral sequence reconstruction,” Nat. Catal., vol. 1, no. 11, pp. 878–888, 2018.

[9] M. Alcalde, “When directed evolution met ancestral enzyme resurrection,” Microb. Biotechnol., vol. 10, no. 1, pp. 22–24, 2017.

[10] B. J. Gomez-Fernandez et al., “Directed -in vitro-evolution of Precambrian and extant Rubiscos,” Sci. Rep., vol. 8, no. 1, pp. 1–11, 2018.

[11] G. K. Hochberg and J. W. Thornton, “Reconstructing ancient proteins to understand the causes of structure and function,” Annu. Rev. Biophys., vol. 46, pp. 247–269, 2017.

[12] J. Jumper et al., “Highly accurate protein structure prediction with AlphaFold,” Nature, vol. 596, no. 7873, pp. 583–589, 2021.

[13] A. Perrakis and T. K. Sixma, “AI revolutions in biology: Protein structure prediction,” EMBO Rep., vol. 22, no. 10, p. e54046, 2021.

[14] J. Yang et al., “Improved protein structure prediction using predicted interresidue orientations,” Proc. Natl. Acad. Sci., vol. 117, no. 3, pp. 1496–1503, 2020.

[15] D. Eramian, “The impact of AlphaFold on protein science: A review,” J. Mol. Biol., vol. 434, no. 5, p. 167630, 2022.

[16] K. Ogami et al., “Protein complex modeling using ancestral sequence reconstruction and AlphaFold2 for elucidating molecular evolution,” Sci. Rep., vol. 12, no. 1, pp. 1– 11, 2022.

[17] E. F. Pettersen et al., “UCSF ChimeraX: Structure visualization for researchers, educators, and developers,” Protein Sci., vol. 30, no. 1, pp. 70–82, 2021.

[18] E. C. Meng et al., “Tools for integrated sequence-structure analysis with UCSF Chimera,” BMC Bioinformatics, vol. 7, p. 339, 2006.

[19] “The UniProt Consortium, “UniProt: the universal protein knowledgebase in 2021,” Nucleic Acids Res., vol. 49, no. D1, pp. D480–D489, 2021.

[20] S. Kuraku, C. M. Zmasek, O. Nishimura, and K. Katoh, “aLeaves facilitates on-demand exploration of metazoan gene family trees on MAFFT sequence alignment server with enhanced interactivity,” Nucleic Acids Res., vol. 41, no. W1, pp. W22–W28, 2013.

[21] L. T. Nguyen, H. A. Schmidt, A. Von Haeseler, and B. Q. Minh, “IQ-TREE: a fast and effective stochastic algorithm for estimating maximum-likelihood phylogenies,” Mol. Biol. Evol., vol. 32, no. 1, pp. 268–274, 2015.

[22] S.Q. Le and O. Gascuel, “An improved general amino acid replacement matrix,” Mol. Biol. Evol., vol. 25, no. 7, pp. 1307–1320, 2008.

[23] J. Sukumaran and M. T. Holder, “DendroPy: a Python library for phylogenetic computing,” Bioinformatics, vol. 26, no. 12, pp. 1569–1571, 2010.

[24] L. McInnes, J. Healy, and J. Melville, “Umap: Uniform manifold approximation and projection for dimension reduction,” arXiv [stat.ML], 2018.

[25] P. J. Cock et al., “Biopython: freely available Python tools for computational molecular biology and bioinformatics,” Bioinformatics, vol. 25, no. 11, pp. 1422–1423, 2009.

[26] M. Mirdita, S. Ovchinnikov, and M. Steinegger, “ColabFold - Making protein folding accessible to all,” bioRxiv, 2021.

[27] H. Ogata et al., “KEGG: Kyoto encyclopedia of genes and genomes,” Nucleic Acids Res., vol. 27, no. 1, pp. 29–34, 1999.

[28] P. Radivojac et al., “A large-scale evaluation of computational protein function prediction,” Nat. Methods, vol. 10, no. 3, pp. 221–227, 2013.

[29] K. Tunyasuvunakool et al., “Highly accurate protein structure prediction for the human proteome,” Nature, vol. 596, no. 7873, pp. 590–596, 2021.

[30] P. Bryant, G. Pozzati, and A. Elofsson, “Improved prediction of protein-protein interactions using AlphaFold2,” Nat. Commun., vol. 13, no. 1, pp. 1–11, 2022.

